# Corrigendum to: T cell apoptosis at the maternal-fetal interface in early human pregnancy, involvement of galectin-1.- A correction with an important twist

**DOI:** 10.1101/374322

**Authors:** Hernan Kopcow

## Abstract

We previously published a paper reporting the apoptotic nature of first trimester human decidual T cells (Kopcow et al 2008). New experiments and data re-analysis produced confirmatory, negative and contradictory results. We had previously shown by TUNEL and sub diploid DNA content analysis (SD) that ex-vivo decidual T cells, isolated either as CD3+ cells or as CD4+/8+ cells, are apoptotic and that the level of apoptosis differs significantly from peripheral blood T cells isolated in a similar way. Data re-examination including data points previously unintentionally ignored results in indetermination in the dataset structure with contradictory statistical significance results between TUNEL and SD and the two isolation approaches. Furthermore, data disaggregation and granular examination reveals discrepant TUNEL and SD readings within individual samples. We have previously shown that decidual CD3+ lymphocytes stain brighter with AnnexinV -Fitc than peripheral blood T cells and other decidual lympohocyte populations. Independent repetitions with modified and alternative isolation protocols using different flow cytometry equipment in the hands of another researcher failed to reproduce those results, while side by side repetition under the published conditions as well as under the alternative isolation protocol reproduced the published pattern when samples were ran in the same equipment used for the publication. A reflection on these results proposes to understand the scientific paper as one among other possible constructed narratives.

## Introduction

In the above cited paper we reported the apoptotic nature of human decidual T cells. Reexamination of the data and new data gathered after publication does not allow to establish with certainty whether decidual T cell (dT) are or are non-apoptotic.

Evidence of dT apoptosis in the paper was based on three observations.

1. Annexin V staining of decidual lymphocytes resulting in a CD3+ lymphocyte population that stained brighter than other decidual lymphocytes with Annexin V-FITC (Fig 4A and A’ in the paper).
2. Flow cytometry TUNEL (Fig 4B and B’ in the paper) and Subdiploid DNA content analysis (SD) (Fig 4C and C’ in the paper) of FACS sorted dT and pT cells.
3. TUNEL and anti-CD3 staining co-localization in serial sections of decidual histological prepartions (Fig 5 in the paper).

Due to reports of difficulties in reproducing the Annexin V staining results.a side by side repetition was conducted and TUNEL and SD data were reexamined

## Results

### TUNEL and subdiploid DNA content analysis

Based on the N reported for each experimental group and the total number of datafiles available, some samples were inadvertenly ommitted from the analysis and one reading was counted twice. A new analysis is presented here including all samples taking into account the following considerations:

- Data disagregation reveals ambiguities in the dataset structure:
- Samples were ran on a FACScalibur flow cytometer on two different days. Most, <u>**but not all**</u>, samples of the group ran on the second day presented a shift to the right of the TUNEL main peak. It is reasonably commonly assumed that TUNEL and SD data for each sample should be somewhat consistent as both are indicative of the level of DNA fragmentation and are obtained from the same experimental tube on a single run. In our dataset TUNEL and SD content values are more consistent if one corrects for the TUNEL peak shift on the second day group.
- The two groups of samples were stored for different times in 70% ETOH at −20C after fixation and before completing TUNEL processing and running on the FACSCalibur flow cytometer. The first group of samples was stored for 2 to 7 days, the second group for 21-30 days. According to the TUNEL KIT manufacturer’s instructions (APO-BRDU™ Kit, BD Biosciences) samples can be kept at −20 for several months, however it can not be ruled out that the different storage time or some other unidentified factor may have affected results, in particular the major TUNEL peak shift to the right of the majority of the second group of samples.

Taking all these facts into consideration, reanalysis of the data including all samples and correcting for the second group TUNEL peak shift, reveals a higher percentage of apoptotic decidual T cells (dT) than peripheral blood T cells (pT) as reported. However, TUNEL differences were statistically significant for cells isolated as CD4/8+ cells but not for cells isolated as CD3+ cells. Conversely SD differences were statistically significant for cells isolated as CD3+ cells but not for cells isolated as CD4/8+ cells (Figure 1-Paper New Figure 4 B’ and C’).

**Figure 1.**
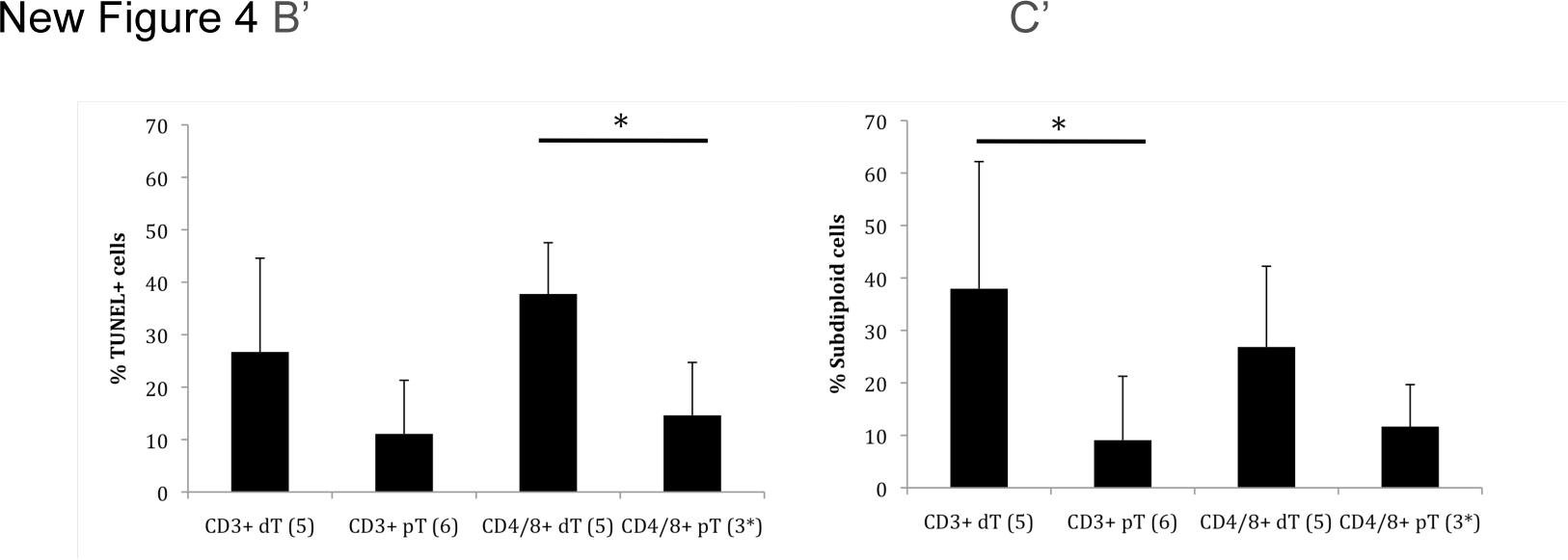
**Paper New Figure 4 B’ and C’** including all samples. Percentage of TUNEL (B’) and subdiploid (C’) decidual T cells isolated as CD3+ or CD4/8+ cells. * p-value <0.05 by one tail non-paremetric T-test. Statistics were only calculated for comparisons of pT and dT samples isolated in a similar way (i.e as CD3+ or as CD4/8+). N of each group is shown in parenthesis.

Therefore the statistical significance analysis results are contrasting between TUNEL and SD and the two cell isolation approaches (ie. as CD3+ or CD4/8+ cells).

Granular examination of the data reveals a data anomaly, a decidual T cell sample (S1, depicted in red Supl Fig 1) ran on the first day, and therefore not affected by the second day TUNEL peak shift, that had a noisy FSC / SSC plot for cells isolated as CD3+ cells, presented discrepant TUNEL and SD values within both isolation approaches. TUNEL values were low and SD values high when cells were isolated as CD4+/CD8+ cells, and TUNEL values were high and SD values low for cells isolated as CD3+ cells. While another dT sample sample (7 o 9w) isolated as CD3+ cells presented high TUNEL and low SD values. Thus these two samples and the TUNEL and SD statistical analysis results considering all samples indicates that the assumption of coherence between different techniques that measure a similar phenomenon is not always valid.

Interestingly, previous attempts by others to establish the apoptotic nature of dT cells have also produced contrasting results between two different techniques, SD flow cytometry and DNA fragmentation analyzed by gel electrophoresis (Jerzak M et al AJRI 1998).

Raw flow cytometry data and flow cytometry histograms used for the analysis above are provided as Supplemental Files 1 and 2. Analysis was done using the Mac version of FlowJo 10.4.2 from Treestar.

### Annexin V staining of decidual lymphocytes-Declining effect and indetermination

The published dT Annexin V - fitc staining results were robust and highly reproducible in the hands of at least two lab members previous to publication. While there was some variability, the vast majority of samples presented a profile similar to the one shown in the paper’s Figure 4 A, with CD3+ decidual lymphocytes (dLs) staining brighter for Annexin V than most other dLs. See examples in Suplemental File 3.

That the brighter annexin V staining of dT cells corresponds to apoptosis and not to cell autofluorescence is supported by preliminary experiments showing in-vitro induction of apoptosis on CD3-dLs by Galectin-1. Exposure of dLs to increasing concentrations of Galectin-1 does not affect the level of Annexin V staining of CD3+ dLs but augments the Annexin V staining Intensity of CD3-dLs up to a fluorescence intensity similar to that of CD3+ dLs. In contrast both CD3+ and CD3-PBLs were negative for Annexin V staining at all galectin 1 concentrations (Fig 2).

**Figure 2.**
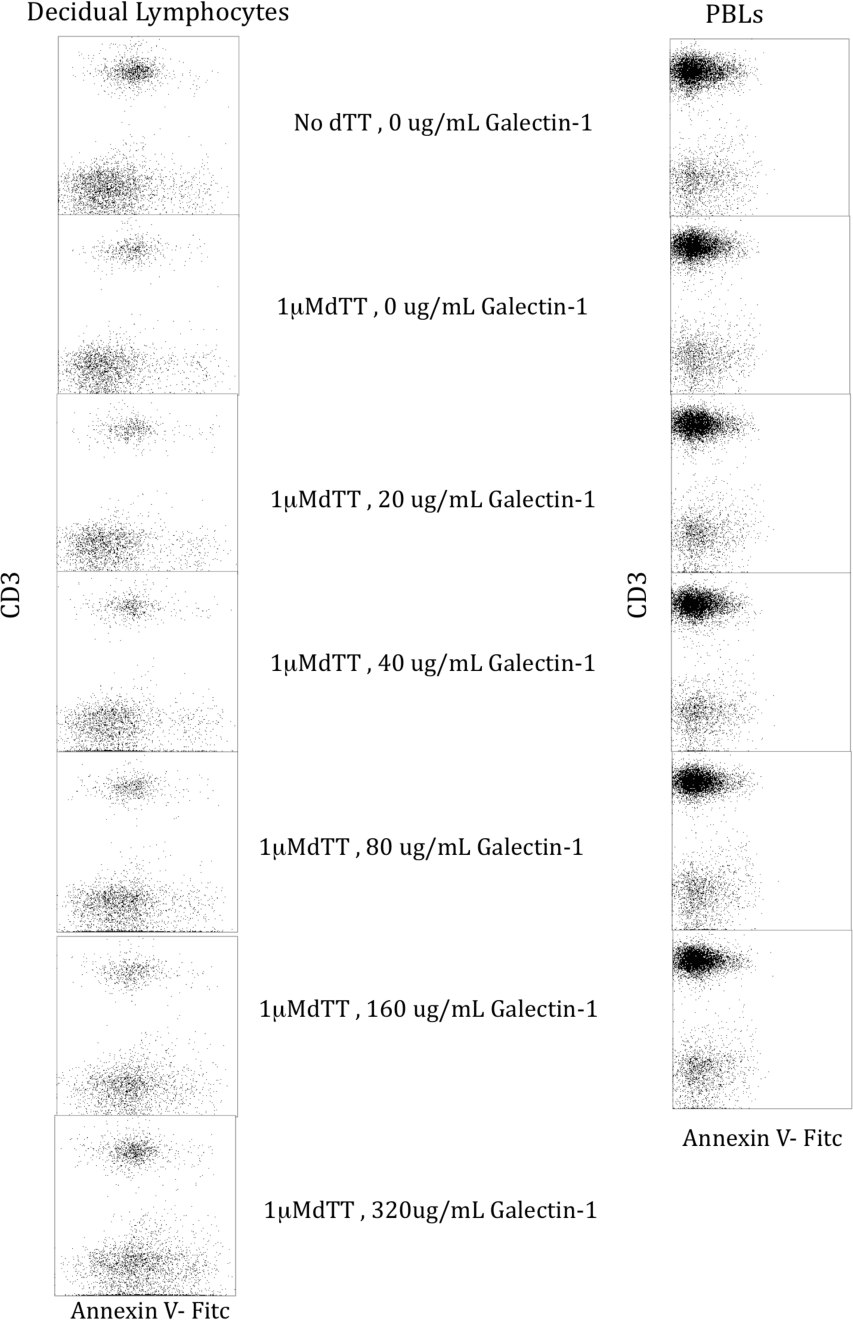
Annexin V-FitC staining profile of decidual Lymhocytes(left) and PBLs (right) exposed for 30 minutes to increasing concentrations of galectin-1 in the presence of 1μM dTT, A control in the absence of galectin-1 and dTT is shown on the top. (n=1)

Therefore ex vivo dT cells were positive for annexin V staining presenting discrepant results for TUNEL and SD depending on the isolation method (as CD3+ or CD4/8+).

### dT Annexin V staining reproducibility problems and effect of incubation at 37 C during isolation

Some time after publication of the paper, there were claims of difiiculty in reproducing the Annexin V staining pattern of dLs isolated using two protocols: The ficol protocol described in the publication which involved DNAse and Collagenase digestion of decidual tissue, obtention of a single cell suspension passing the cells through 100um, 70um, and 40um cell strainers, Incubation at 37C for 90 to 120min to remove adherent cells, and subsequent lymphocyte isolation by ficol (protocol described in Kopcow et al 2008). This isolation procedure is referred from here on as the Ficol protocol. The second protocol referred as the Percol protocol, was similar but omitted the incubation at 37C and achieved lymphocyte isolation using a 20%-45%-68% Percol gradient rather than Ficol and is described elsewhere (Tilburgs et al 2015). Those claims were taken with great surprise given the robustness of the dLs annexin V staining pattern previous to publication.

As one of the differences between the two protocols was the incubation at 37 C, the effect of 37 C incubation during the ficol isolation protocol on dLs AnnexinV staining patterns was evaluated focusing on CD3+ dT and decidual NK cells (dNK). In two out of three samples Annexin V-Fitc staining of dT cells augmented after 90 minutes incubation at 37 C. The third sample, however, presented dT cells with high levels of annexin V stainig even without incubation at 37C. dNK cell annexin V staining pattern also changed, dNK cells being heterogeneous for annexin V staining before incubation at 37 C and changing to two populations, a major Annexin V negative population and a very minor annexin V super bright population after incubation (Figure 3). Whether these patterns are the result of apoptosis induction during the incubation at 37 C or the removal of dead cells by ficol, or in the case of dNK cells reversion from apoptosis deserves further investigation.

**Figure 3.**
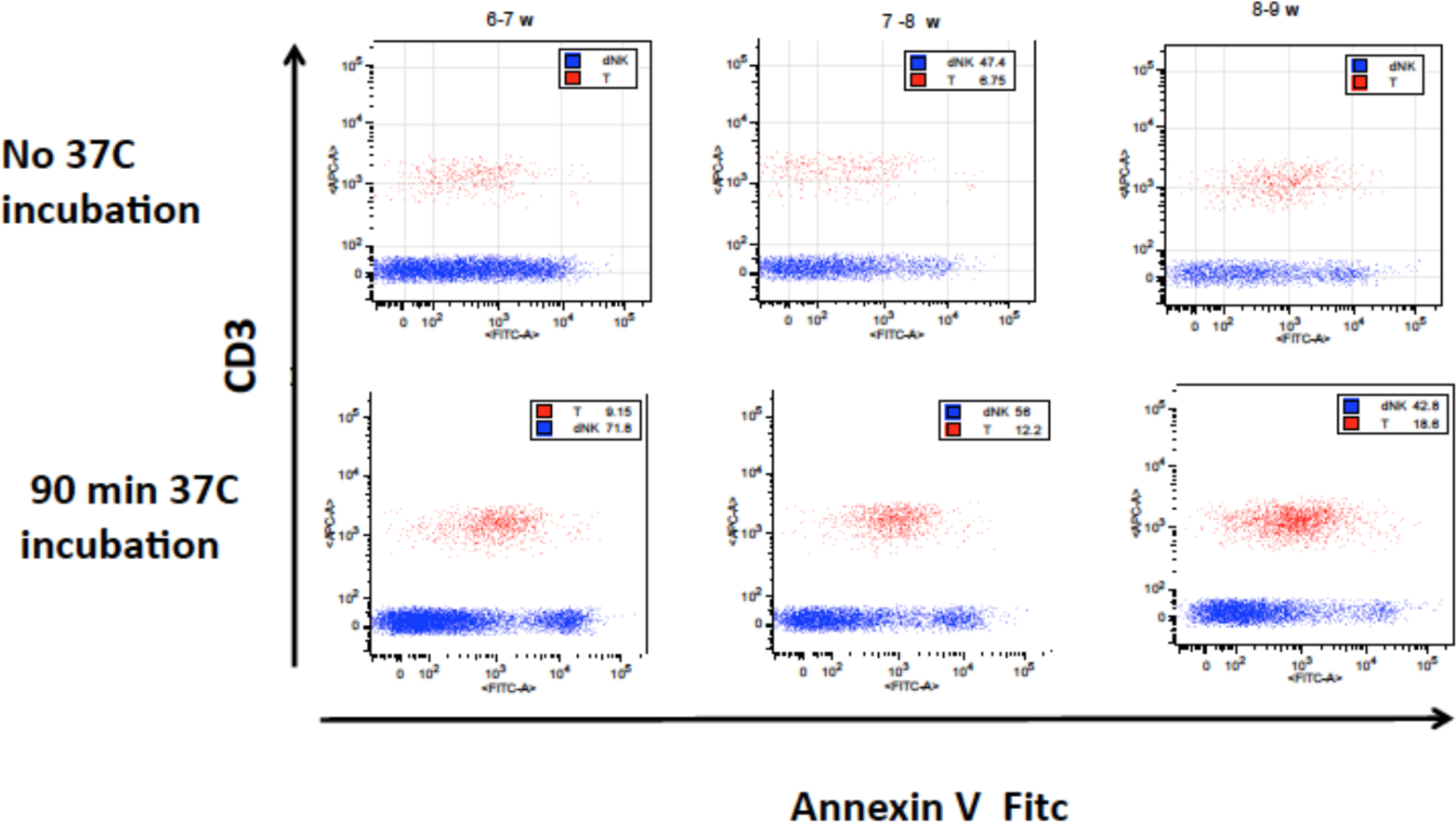
Effect of 37C incubation on Annexin V stianing of decludial Lymphocytes in the Ficol isolation protocol. Annexin V staining profile of dLs gated on CD3+ dT (Shown in red) cells and CD56+ dNK cells(shown in blue) in three decidual lympohocyte samples isolated by the Ficol isolation protocol including (bottom panels) or not including (top pannels) a 90 minutes 37C incubation step. Samples were processed on the same day they were collected. Plots were generated on a FACSCanto flow cytometer using FACS DIVA software and analyzed. Plots are gated on a lymphocyte FSC/SSC gate, 7AAD-cells, and either CD3+ (red dots) or CD56+ (blue) cells. Gestational age of each decidual sample is shown above the plots on the top.

**Figure 4.**
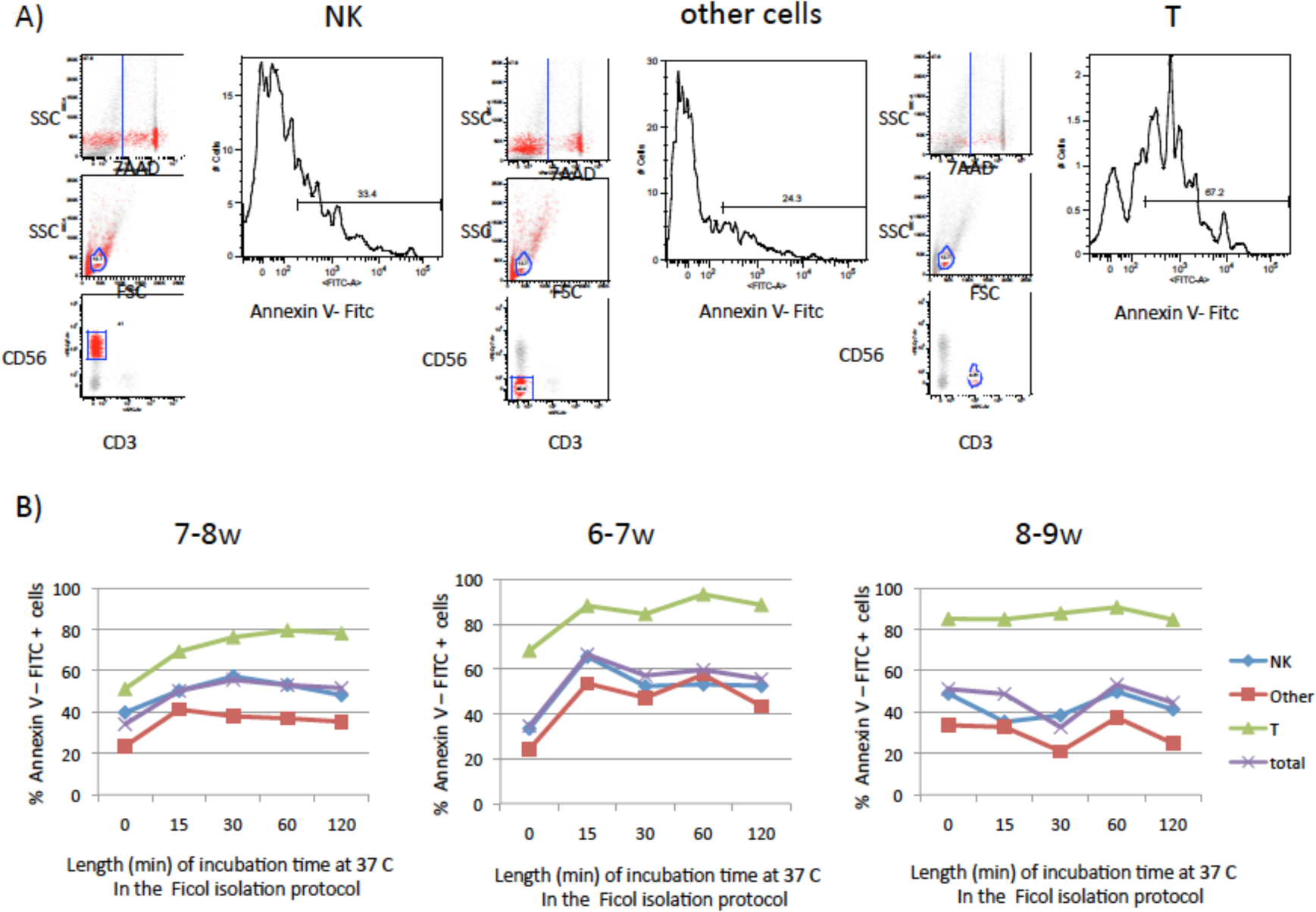
Time course analysis of the effect of incubation at 37C in the ficol protocol on the annexin V staining profile of decidual lymphocytes. A) Gating strategy to quantify the percentage of annexin V-Fitc positive cells among decidual NK cells (left), decidual T cells(right), and other decidual lymphocytes (center). B) % of Annexin V-Fitc + cells at different incubation times at 37 C during the Ficol Isolation protocol. Results on three decidual samples is shown. The isolation procedure was initiated a day after collecting decidual tissue samples. Estimated gestational age is indicated above the graph of each sample. Decidual NK cells, decidual T cells, other cells, and total lympocytes are shown in each graph

A time course analysis on the same samples in a cell isolation performed the following day revealed that in two of the samples the increase in dT annexin V staining took place on the first 15 to 30 minutes of the 37C incubation. However the third sample was positive before incubation and remained at about the same level of staining throughout the 120 minutes at 37 C (Fgure 4)

Therefore while two samples supported the idea that incubation at 37C affects the level of annexin V staining on dT cells, a third sample presented dT cells that stained bright for annexin V independently of the incubation at 37 C thus introducing variability, an element of indetermination or an anomaly depending on how it is preferred to be interpreted.

### Side by side repetition of dT Annexin V staining with Ficol and Percol protocols

Careful examination of the Ficol protocol used in the repetition that had failed to reproduce the published Annexin V staining pattern revealed the following differences with the published protocol: In the failed repetition attempt the 37C incubation and Ficol isolation steps had been swapped. Annexin V staining was done on ice instead of at room temperature. In the publication samples were ran on a FACScalibur flow cytometer using CellQuest software, in the failed attempts on a LSRII flow cytomoter with FACS Diva software.

Experiments were then repeated side by side using both protocols, Ficol as published and Percol, running samples both on the Facsclibur flow cytometer and on an LSRII flow cytomoter. Details of the protocols used and their differences are provided as screenshots of notes taken after the repetitions (Supplemental file 4).

Results in the Facscalibur flow cytometer reproduced the published pattern of annexin V+ CD3+ dLs for cells isolated with the Ficol protocol, and according to notes, as well as with the Percol protocol(Supplemental file 4). it would be important to confirm these results with the actual data. No attention was given to the LSRII results as under the published conditions samples ran on the FACSCalibur flow cytometer confirmed the published pattern.

A few weeks later, however, the person that originally challenged the published results obtained, in an independent repetition, negative stainings with both protocols using a LSRII flow cytometer with FACS DIVA software. Interestingly, in my own hands (HK) a later isolation and evaluation using a FACSCanto flow cytomoter with FACSDiva Software also presented an Annexin V negative staining pattern on CD3+ cells, but this was an anomaly as in other isolations cells maintained the published pattern.

### TUNEL and anti-CD3 staining co-localization in serial sections of decidual histological prepartions

The published colocalization of anti CD3 and TUNEL staining in serial sections of histological preparations support the idea that decidual T cells are apoptotic, however because staining for each marker was performed on serial sections the results are supportive but not conclusive, as it can not be ruled out the possibility that TUNEL staining corresponds to cells adjacent to CD3+ cells instead of CD3+ cells themselves.

## Discussion

Science assumes coherence between techniques that measure similar things, and aggregates data in the form of statistical analysis. Such aggregation some times masks data anomalies that challenge the assumption of coherence.

For instance the TUNEL and SD analsyisis presented here reveals contrasting statistical significance results between the two techniques and the way the cells were isolated, either as CD3+ or as CD4+/8+ cells. The conclusion to which one would arrive would be different if only one technique had been performed. Similarly particular samples presented discrepant SD and TUNEL results while both techniques evaluate the extent of DNA fragmentation on the same physical tube and in a single run.

Annexin V staining of decidual lymphocytes isolated with the Ficol protocol presented robust highly reproducible results before publication, however later challenges to the reproducibility of the results support the idea of declining effect, that reproducibility declines over time with the number of repetitions (Schooler 2011). A number of differences could be pointed out between the published results and the failed repetition attempts, including the temperature at which the annexin V staining was performed (Room temperature or on ice) Flow cytometer and software used, swaping of the Ficol and 37C incubation steps, the researcher performing the experiment, or other unidentified causes. Thus results can be sensitive to techniques and protocol variations or the person performing the experiment.

Annexin V staining patterns of some samples, but not all samples, were sensitive to the incubation at 37C. This fact could make one think that the Percol protocol could be more appropriate as it does not have a 37C incubation step. All previous experiments using the Percol protocol that had failed to show Annexin V+ dT cells were ran on a LSR II flow cytomoter. However when cells isolated by the Percol protocol were ran on the FACScalibur flow cytometer in the side by side repetition they presented a staining pattern with Annexin V positive cells similar to the published one, according to notes taken at that time. It would be important to be able to confirm these results with the actual data. Furthermore, as the samples were also ran on the LSRII flow cytometer, and the Percol protocol was always performed by the same person, comparison of the Percol-LSRII results and the Percol-FAcscalibur results, would be instrumantal on establishing if there is an effect introduced by the flow cytometry platform, or there was some other factor introducing a day to day variability. If the results were similar to the FAcscalibur results, that would point out to inter day variability, or ambiguity.

If the Percol protocol would be considered more advantageous, it would be necessary to reexamine some of the most cited papers in dNK cell biology that used the Ficol isolation protocol (Koopman et al 2003 for instance), and ultimetly in the face of discrepant results, or discrepant gene expression profiles as is the case of the cited paper, one would be forced to either, choose for one or the other, or accept both possible options as valid, accepting again the existence of indetermination in the experimental setting.

In conclusion in the research process one makes choices, as selecting one or another protocol, or technique, that may affect the observed results. Here a mistake led to the publication of coherent Annexin V, TUNEL and SD data supporting the apoptotic nature of dT cells. Relization and correction of that mistake introduces discrepancy between the Annexin V results, and TUNEL and SD results, the last two being contrasting one with the other. Furthermore, the robust Annexin V staning results were challenged with cells isolated with the alternative Percol protocol, but at the side by side repetition, even the Percol protocol confirmed the published results according to notes. Thus, it is often the case that the requirement for certainty and coherence of the publication process, sometimes demanding the exclusion of negative data during the review process, ends up leading to the construction of a narrative, among other possible narratives. Given that our minds need of linear narratives in order to make thing intelligible, this is acceptable, as long as it is understood as such, leaving open the possibility for alternative experimental and interpretative perspectives.

